# Orthologous synteny provides robust structural evidence for the ancestral angiosperm ε-WGD

**DOI:** 10.64898/2026.03.05.709955

**Authors:** Ren-Gang Zhang, Martin A. Lysak, Hong-Yun Shang, Yuan-Nian Jiao, Yong-Peng Ma

## Abstract

The ancestral angiosperm whole-genome duplication (ε-WGD) is a pivotal yet contentious event that has long been debated. A recent study refuted its occurrence based on deviations from expected retention patterns of dosage-sensitive genes. Here, we provide both structural (orthologous synteny) and phylogenetic (subgenome-aware) evidence to revisit this debate. With improved annotation of the *Ginkgo* genome, we uncover a clear 1:2 orthologous synteny ratio between *Ginkgo* and early-diverging angiosperms (*Amborella* and *Aristolochia*), which lack lineage-specific WGDs. Subgenome-aware phylogenomic analyses further demonstrate that the *Amborella*/*Aristolochia* subgenomes diverged after the gymnosperm–angiosperm split but before the radiation of extant angiosperms. Together, these findings provide direct structural evidence for an ancestral angiosperm tetraploidization in the early Jurassic, following divergence from gymnosperms. We argue that the recent refutation of the ε-WGD relies on restrictive assumptions regarding ploidy level and duplicate retention that may not hold for deep-time polyploidy events. Our results provide a robust resolution to this long-standing debate and reaffirm the ε-WGD as a shared synapomorphy of extant angiosperms.

## Introduction

Ancient whole-genome duplications (WGDs) have been inferred at the dawn of seed plants (ζ-WGD, ∼319 million years ago) and angiosperms (ε-WGD, ∼192 million years ago)^1^. However, the occurrence of these WGDs has remained a subject of long-standing debate^2^, hindering our understanding of how polyploidy contributed to the evolutionary success of seed and flowering plants. Recently, Shi and Van de Peer (2026) validated the ζ-WGD but refuted the ε-WGD based on the retention patterns of dosage-sensitive gene duplicates^3^. They argued that the duplication signals mapped to the ancestral angiosperm branch were inconsistent with expectations for an ε-WGD scenario. While their study provides a novel statistical perspective, we interpret both their data and the broader genomic evidence differently.

In this study, we revisit the ε-WGD using state-of-the-art orthologous synteny analyses and subgenome-aware phylogenomics. Our results provide robust structural and evolutitionary evidence supporting the occurrence of the ε-WGD, providing the solid evidence needed to settle this long-standing debate.

## Results and Discussion

We first used orthologous synteny analysis^4^ to systematically compare gymnosperm and angiosperm genomes. To provide a high-quality gymnosperm reference, we re-annotated the genome of *Ginkgo biloba*^5^, an early-diverging gymnosperm, increasing its BUSCO completeness from 63.5% to 93.1%. Despite syntenic fragmentation, orthologous synteny between *Ginkgo biloba* and *Amborella trichopoda* — the earliest-diverging angiosperm without lineage-specific WGDs — reveals a clear and consistent 1:2 correspondence (**Fig. 1A**). Specifically, one genomic region in *Ginkgo* aligns with two distinct orthologous regions in *Amborella*. This genome-wide 1:2 syntenic pattern provides direct evidence that *Amborella* underwent a tetraploidization event after its divergence from gymnosperms, corresponding to the ancestral angiosperm ε-WGD. This orthologous 1:2 signal remains robust across additional angiosperms. Species lacking post-angiosperm WGDs, such as *Aristolochia*, retain the 1:2 pattern, whereas lineages with a subsequent WGD exhibit higher ratios (e.g., 1:4 in *Warburgia*), consistent with their independent polyploid histories (**Fig. 1B–C, Fig. S1–S2**).

**Figure 1.**
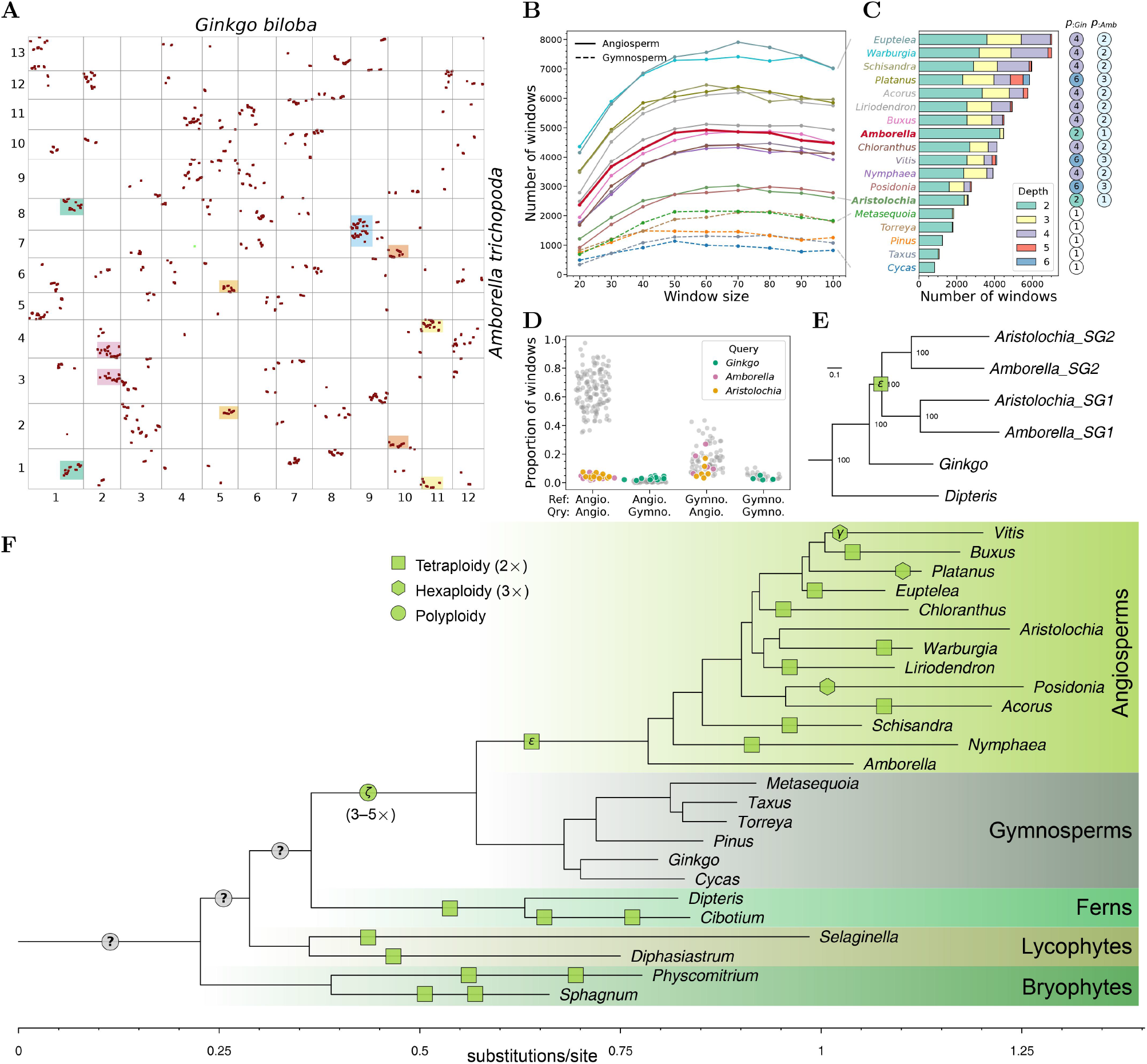
Orthologous synteny and phylogenomic evidence for the ancestral angiosperm ε-WGD. **(A)** Dot plot showing orthologous synteny between *Ginkgo biloba* and *Amborella trichopoda*. Colored shading highlights multiple examples of 1:2 orthologous synteny. **(B)** Abundance of genomic windows with orthologous syntenic depth ≥ 2 relative to the *Ginkgo* reference genome. Values were calculated using a sliding window approach (window size = 20--100 genes, step size = 1 gene) across the *Ginkgo* genome for each representative angiosperm and gymnosperm genome analyzed. **(C)** Composition of orthologous syntenic depth (depth ≥ 2) in gymnosperm and angiosperm genomes relative to the *Ginkgo* reference (window size = 50 genes, step size = 1 gene). Numbers in circles on the right indicate inferred ploidy relative to *Ginkgo* (*p*_:*Gin*_) and *Amborella* (*p*_:*Amb*_) based on syntenic depth patterns (see also evidence in Fig. S1--S3). **(D)** Proportions of genomic windows exhibiting orthologous syntenic depth ≥ 2 in pairwise comparisons between angiosperm and gymnosperm genomes (window size = 50 genes, step size = 1 gene across each reference genome). **(E)** Phylogenetic reconstruction of the relationships among the two *Amborella*/*Aristolochia* subgenomes (SG1 and SG2) and *Ginkgo*. The tree was rooted using *Physcomitrium* and *Selaginella* (not shown). Numbers at the nodes represent bootstrap support values based on 1,000 replicates. **(F)** Phylogenetic framework of land plants illustrating the inferred positions of WGD events. The tree was rooted using streptophyte algae (not shown). Different shapes along branches represent polyploidization events with different inferred ploidy levels (circles denote events with undetermined ploidy). Green-filled shapes indicate WGDs supported by syntenic evidence, while grey-filled shapes denote speculative inferences based on retention patterns of dosage-sensitive gene duplicates reported by Shi and Van de Peer (2026). Placement of symbols along branches is approximate and intended for illustrative purposes only.

In striking contrast, comparisons between *Ginkgo* and selected gymnosperm genomes consistently show a 1:1 syntenic ratio (**Fig. 1B--C, Fig. S3A**). These gymnosperms also generally exhibit 1:2 or higher ratios when compared with angiosperms (**Fig. 1D, Fig. S3B**), thereby restricting the WGD signal to the angiosperm lineage. This reciprocal pattern supports placement of the ε-WGD in the common ancestor of all extant angiosperms. The presence of paralogous synteny in *Amborella*, despite its relatively low duplicate retention (**Fig. S4A**), provides additional structural evidence confirming the ε-WGD event.

To further validate the ε-WGD, we partitioned the duplicated syntenic blocks of *Amborella* into two subgenomes and performed a subgenome-aware phylogenomic analysis^6^. The resulting phylogeny consistently recovered the two subgenomes as sister lineages (**Fig. 1E**). Crucially, divergence between these subgenomes occurred after the split between the *Ginkgo* and angiosperm lineages, but before the divergence of *Amborella* and *Aristolochia* (**Fig. 1E**). The concordance between subgenome phylogeny and orthologous synteny patterns provides unequivocal evidence that the doubling of the *Amborella* genome resulted from a shared ancestral angiosperm WGD, rather than from more ancient seed plant duplications. The integration of structural and phylogenomic signals thus establishes the ε-WGD as a foundational event in angiosperm evolution (**Fig. 1F**).

Synteny is widely recognized as the gold standard for polyploidy inference^2,3^, providing more direct and definitive evidence than gene-based methods such as gene tree-species tree reconciliation or *K*_S_distributions. Historically, however extensive studies have emphasized paralogous synteny — internal duplication signals within a single genome — while overlooking the critical value of orthologous synteny. In contrast, orthologous synteny offers a straightforward, reliable, and versatile framework for identifying polyploidy events, inferring the resulting ploidy levels, and placing these events within phylogenetic trees^4^. This comparative approach is particularly powerful when paralogous syntenic signals have been eroded by long-term biased fractionation, as exemplified by the *Amborella* genome (**Fig. S4A**). Importantly, while syntenic signals are expected to erode over vast evolutionary timescales due to massive gene loss and chromosomal rearrangements, we find substantial orthologous syntenic remnants that remain remarkably well preserved between gymnosperms and angiosperms. This deep structural conservation provides a pivotal structural foundation for confidently inferring both the occurrence and the phylogenetic placement of the ε-WGD.

In the analysis by Shi and Van de Peer (2026), a large number of dosage-sensitive gene duplicates were mapped to to the ancestral angiosperm branch^3^. However, they dismissed the ε-WGD primarily because the observed duplication frequency failed to meet their expected 2:1 (or 1:1) ratio of angiosperm-to-seed-plant duplication nodes (original Fig. 5 in^3^). We argue that this expectation rests on two fragile assumptions. First, the proposed 2:1 model assumes that both the seed plant (ζ-WGD) and angiosperm (ε-WGD) polyploidy events were simple tetraploidizations (i.e., a 2-fold increase in ploidy). However, our synteny analysis of the *Ginkgo* genome shows paralogous synteny depths ranging from 2 to 4 (**Fig. S4B**), consistent with a 3-to 5-fold increase in ploidy associated with the ζ-WGD. This pattern may even reflect two sequential WGDs rather than as single event. If the ζ-WGD event represented, for instance, a hexaploidization (3-fold ploidy increase), the expected ratio would shift to 3:3 (duplicated gene pairs) or 3:2 (phylogenetic nodes), rendering their 2:1 expectation mathematically invalid. Second, the authors assume that duplicates derived from a more recent WGD should necessarily exceed those retained from an older WGD within the same lineage. While this can apply to polyploids with ongoing rediploidization, it is unlikely to apply to extremely ancient events in which rediploidization has long been completed. For such deep-time WGDs, the number of retained duplicates is more likely to be governed by the stabilization reached after long-term gene fractionation^7–9^, rather than simple event age. Consequently, it is challenging to predict which ancient WGD should exhibit higher retention. Indeed, given that the origin of seed plants constituted a more fundamental genomic and morphological innovation than the subsequent emergence of angiosperms^10^, it is plausible that more duplicates were retained from the ζ-WGD than from the ε-WGD. Notably, this interpretation is consistent with their own reported higher retention rate (*q*_ζ_) compared to *q*_ε_ (their Fig. 6C in ^3^). Shi and Van de Peer (2026) further invoked the lineage-specific ohnolog resolution (LORe) model as an alternative explanation, proposing that delayed and lineage-specific post-WGD fractionation could cause a shared WGD event to appear as independent WGDs in descendant lineages. However, fundamental to this model is the expectation of a 2:2 orthologous mapping pattern between the daughter clades^11^. In contrast, the 1:2 (or higher) orthologous syntenic ratios we consistently observed between gymnosperms and angiosperms are incompatible with this model. Finally, while the pioneering study by Jiao et al.^1^ relied primarily on gene-tree-based dating — an approach inherently subject to phylogenetic and temporal uncertainty — our synteny-based evidence provides a robust structural confirmation that the ε-WGD indeed represents a shared synapomorphy of all extant flowering plants.

While we disagree with Shi and Van de Peer’s primary conclusion regarding the validity of the ε-WGD, we emphasize that their methodology remains highly valuable. When reinterpreted under a more flexible, ploidy-aware model, their data in fact provide complementary support for ancient polyploidy events, including the ε-WGD. Given that syntenic signals inevitably erode over deep evolutionary time and become nearly undetectable for the very deep ancestral nodes of land plants, Shi and Van de Peer’s approach offers an important alternative for detecting very ancient WGDs. Their results (original Fig. 5B--C in their paper) already hint at potential WGDs at very deep phylogenetic levels, such as in the common ancestor of vascular plants (**Fig. 1F**), where syntenic evidence is largely vestigial. We hope that by resolving the long-standing debate over the ε-WGD through integrated syntenic and subgenome-aware phylogenomic evidence, the field is now positioned to shift its focus toward these even more ancient polyploidy events. Understanding the evolutionary significance of WGDs during the early evolution of land plants is fundamental, as subsequent rediploidization processes likely fueled key innovations and underpinned the remarkable diversification of the terrestrial flora^12–15^.

## Supplementary Information

**Figure S1**. Orthologous synteny between *Amborella trichopoda* and other 12 angiosperm genomes.

**Figure S2**. Orthologous synteny between *Ginkgo biloba* and other 12 angiosperm genomes.

**Figure S3**. Orthologous synteny of *Ginkgo biloba* (**A**) and *Amborella trichopoda* (**B**) against five other gymnosperm genomes.

**Figure S4**. Paralogous synteny within *Amborella trichopoda* (**A**) and *Ginkgo biloba* (**B**). Subplots: **a**) dot plot colored by *K*_S_, **b**) histogram of *K*_S_, using the same color map as the dot plot, **c**–**d**) synteny depth across 50-gene windows (window step = 1 gene).

**Table S1**. Summary of genomes used in this study.

## Methods

### Data collection and genome annotation

To improve the genomic resources for this study, we re-annotated the *Ginkgo biloba* genome^5^, which served as the primary reference genome. A total of 1,017 RNA-seq datasets were retrieved from the Sequence Read Archive (SRA) and the Genome Sequence Archive (GSA). These raw reads were aligned to the *Ginkgo* genome using minimap2 v2.30-r1287^16^, followed by transcript assembly with StringTie v2.1.5^17^. We then employed EviAnn^18^ to generate an evidence-based gene annotation. For protein evidence, we utilized consensus sequences from gymnosperm gene families previously generated^19^. The re-annotation yielded 20,908 high-confidence protein-coding genes, and annotation completeness was assessed using BUSCO v5.3.2^20^ with the embryophyta_odb10 database. The *Amborella* genome annotation was obtained from the recently published chromosome-level version^21^. For other angiosperms, we selected representatives that have undergone at most one additional polyploidy event. For other gymnosperms, we selected representatives from major lineages that have not experienced any additional polyploidization events. Detailed sources for all genomes used in this study are provided in Supplementary Table S1.

### Identification of orthologous synteny and reconstruction of species tree

Orthologs and paralogs were inferred using OrthoFinder v2.3.1 (parameter: -M msa)^22^. Syntenic blocks were identified using WGDI v0.6.2 with the -icl option^23^. Synonymous substitution rates (*K*_S_) for syntenic gene pairs were estimated using the -ks option in WGDI. Finally, orthologous synteny was rigorously identified by integrating orthology and synteny data using SOI v1.3.0^4^, ensuring that the identified blocks represented speciation-derived relationships rather than ancient paralogy. To determine the depth of the orthologous synteny, we employed a sliding window approach across the reference genome. Using the SOI depth function, we quantified the synteny depth as the number of query genomic segments covering each focal window.

A species tree was reconstructed using the SOI phylogenomics pipeline^4^ based on the orthologs identified by OrthoFinder. A concatenated supermatrix was constructed from 1283 single-copy genes (allowing a maximum of 20% missing taxa), and a maximum-likelihood tree was inferred using IQ-TREE v2.2.0.3^24^ with 1,000 bootstrap replicates.

### Inference of paleopolyploidy

Paleopolyploidy events were inferred primarily through genome-wide orthologous synteny patterns, which provide direct evidence^4^. For instance, a 1:1 orthologous synteny depth ratio between *Aristolochia* and *Amborella* indicates that neither lineage has undergone an additional post-divergence WGD. Conversely, a clear 2:1 ratio between *Warburgia* and *Amborella* confirms an independent WGD specific to the *Warburgia* lineage. To further validate these inferences, we analyzed the *K*_S_ distribution of paralogous syntenic gene pairs, as a distinct *K*_S_ peak corresponding to duplicated syntenic blocks would provide independent support for the occurrence of a WGD event.

### Subgenome assignment and phylogenetic reconstruction

For the *Amborella* genome, we manually assigned the duplicated syntenic blocks into two subgenomes with the assistance of the WGDI toolkit, following the guideline of Zhang et al.^6^ with minor modifications. For genomic regions where subgenome phasing was ambiguous across different chromosomes, blocks were assigned to subgenomes randomly (relative phasing); this strategy does not affect the recovery of the sister relationship between the two subgenomes^6^. Next, the subgenomes of other angiosperms were assigned by mapping them to the *Amborella* genome based on the orthologous relationships using WGDI -km option. Syntenic gene lists were generated using the -a option in WGDI. For outgroup species where synteny was no longer conserved, orthologous genes were retrieved to serve as surrogates for syntenic orthologs. Based on the -at option in WGDI, we generated codon-based alignments for each orthologous group. These individual alignments (130 gene loci, at least 6 out of 8 sequences per locus) were subsequently concatenated into a supermatrix. A phylogenomic tree was then reconstructed from this concatenated alignment using IQ-TREE with 1,000 bootstrap replicates.

## Data Availability

The improved gene annotations of *Ginkgo biloba* are deposited in Figshare (https://doi.org/10.6084/m9.figshare.31410387).

## Acknowledgments

We thank Hao Yang for assistance with data download.

## Author contributions

YPM and RGZ conceived and designed the study. RGZ collected and analyzed the data, and drafted the manuscript; YPM, MAL and RGZ revised the manuscript; all authors approved the final manuscript.

