## Supplemental figures for "Orthologous synteny provides robust structural evidence for the ancestral angiosperm ε-WGD"

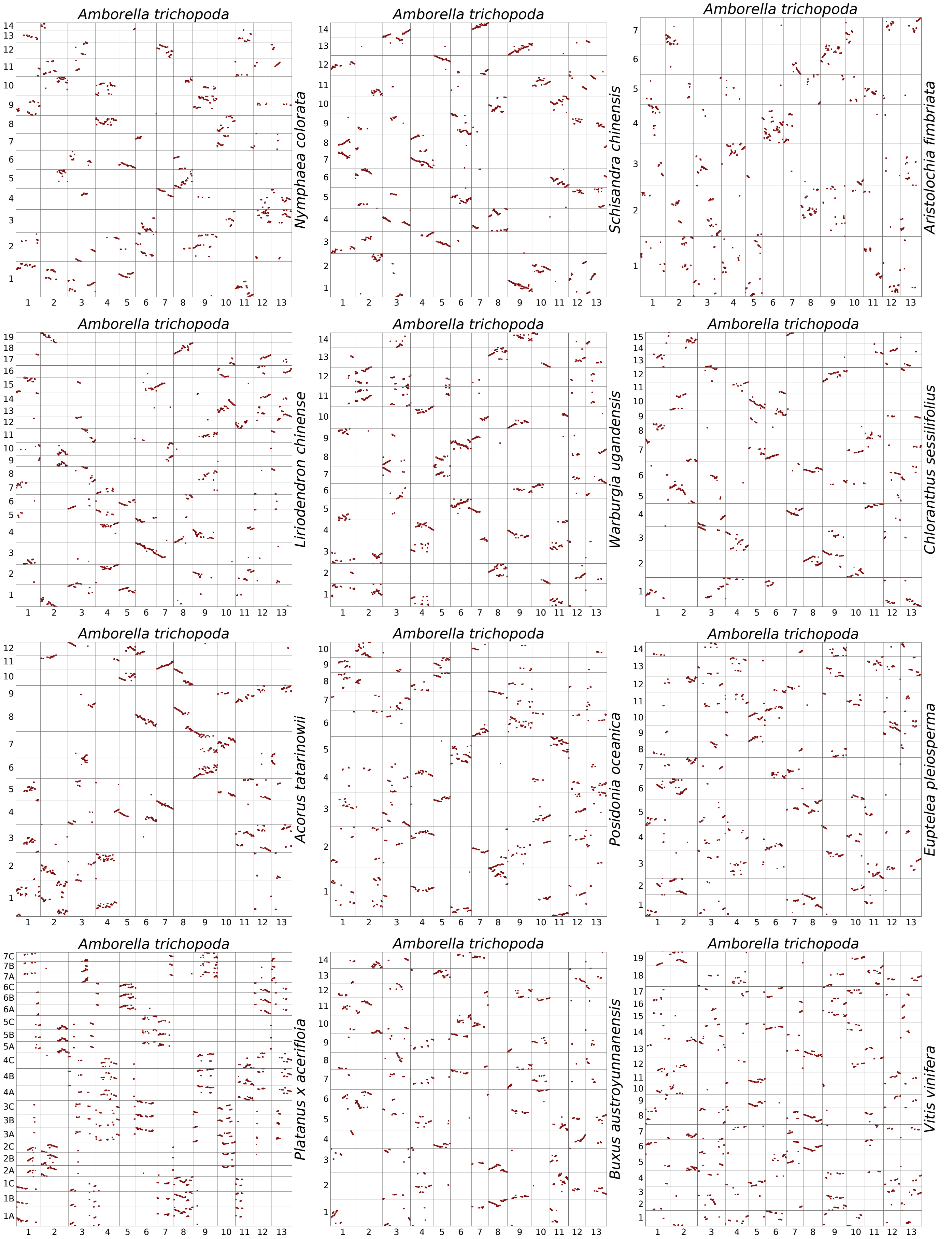


**Supplementary Figure 1. Orthologous synteny between *Amborella trichopoda* and 12 other angiosperm genomes.**

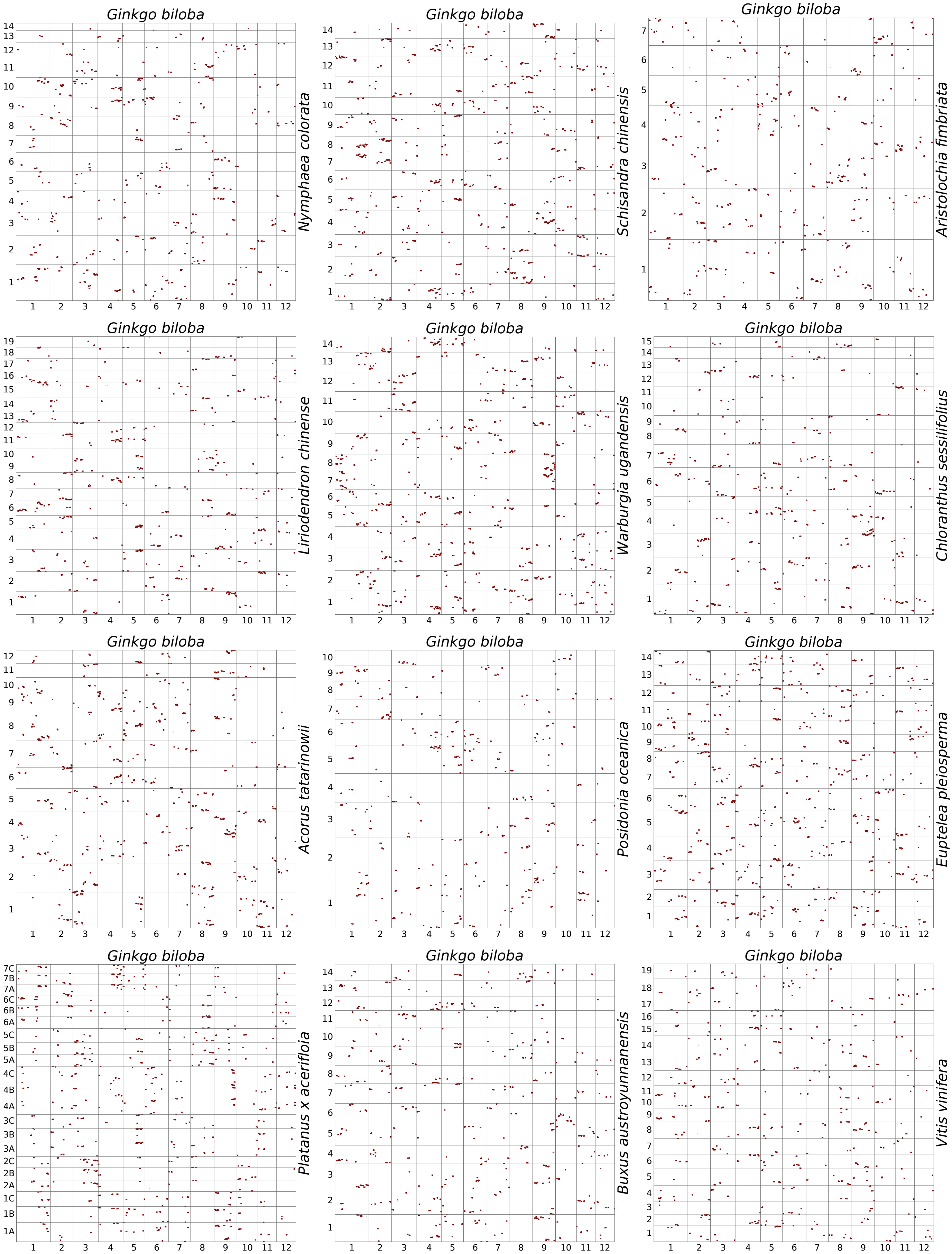


**Supplementary Figure 2. Orthologous synteny between *Ginkgo biloba* and 12 other angiosperm genomes.**

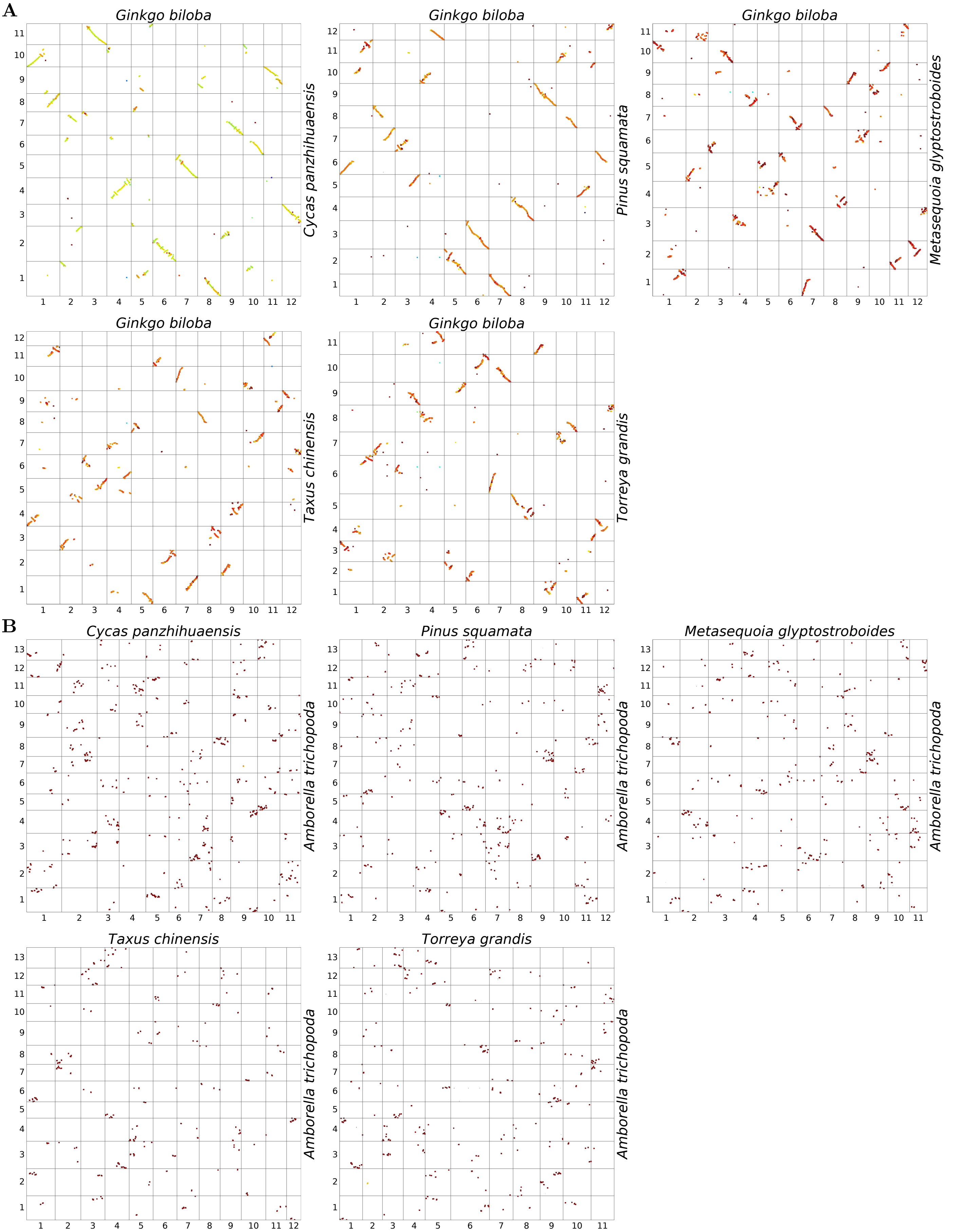


**Supplementary Figure 3. Orthologous synteny of *Ginkgo biloba* (A) and *Amborella trichopoda* (B) against five other gymnosperm genomes.**

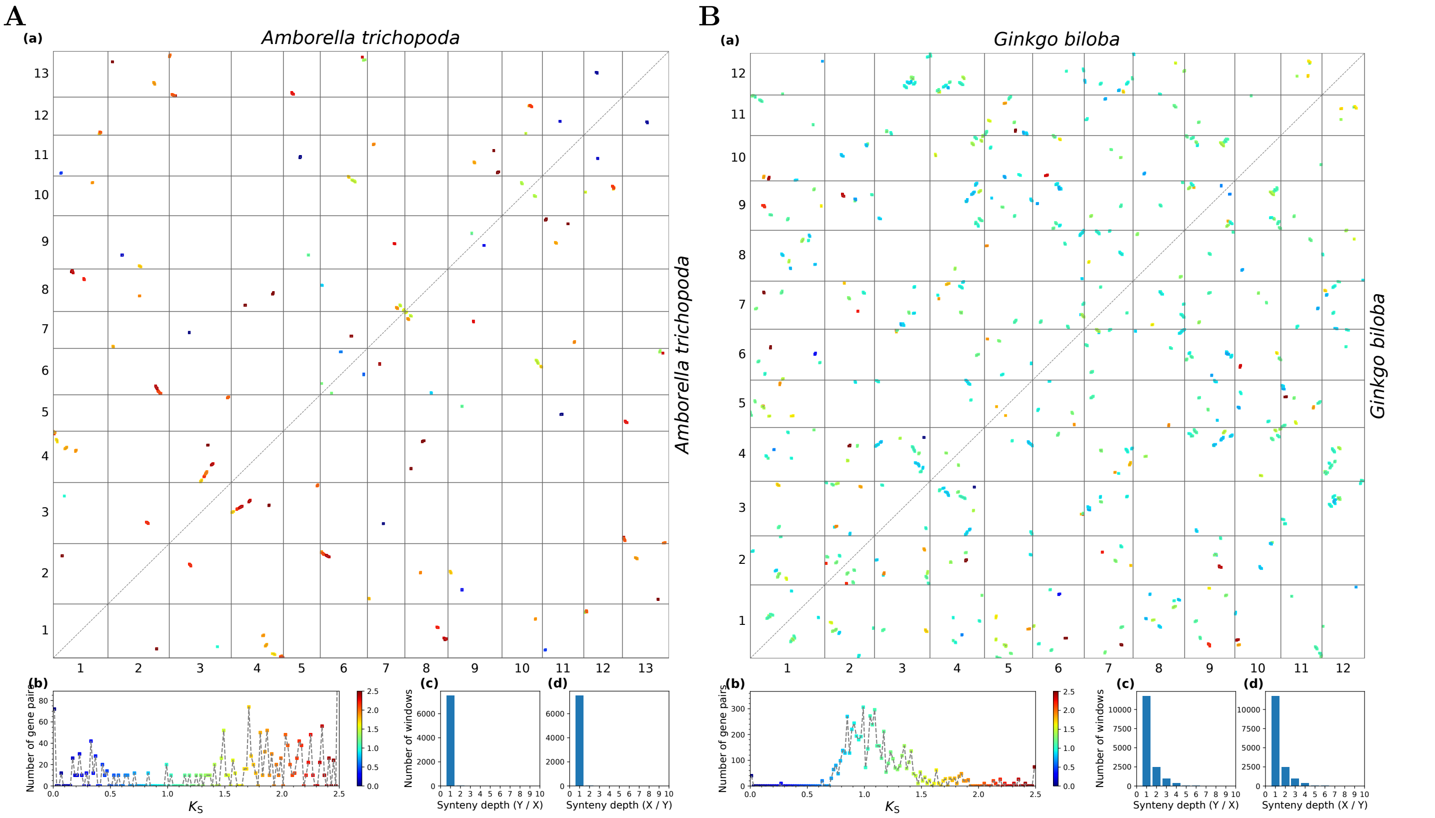


**Supplementary Figure 4. Paralogous synteny within *Amborella trichopoda* (A) and *Ginkgo biloba* (B).** Subplots: a) dot plot colored by $K_S$, b) histogram of $K_S$, using the same color map as the dot plot, c–d) synteny depth across 50-gene windows (window step = 1 gene).
